## Supplementary Note, Supplementary Figure for "Connecting high-resolution 3D chromatin organization with epigenomics"

### Supplementary materials for paper titled "Connecting high-resolution 3D chromatin organization with epigenomics"

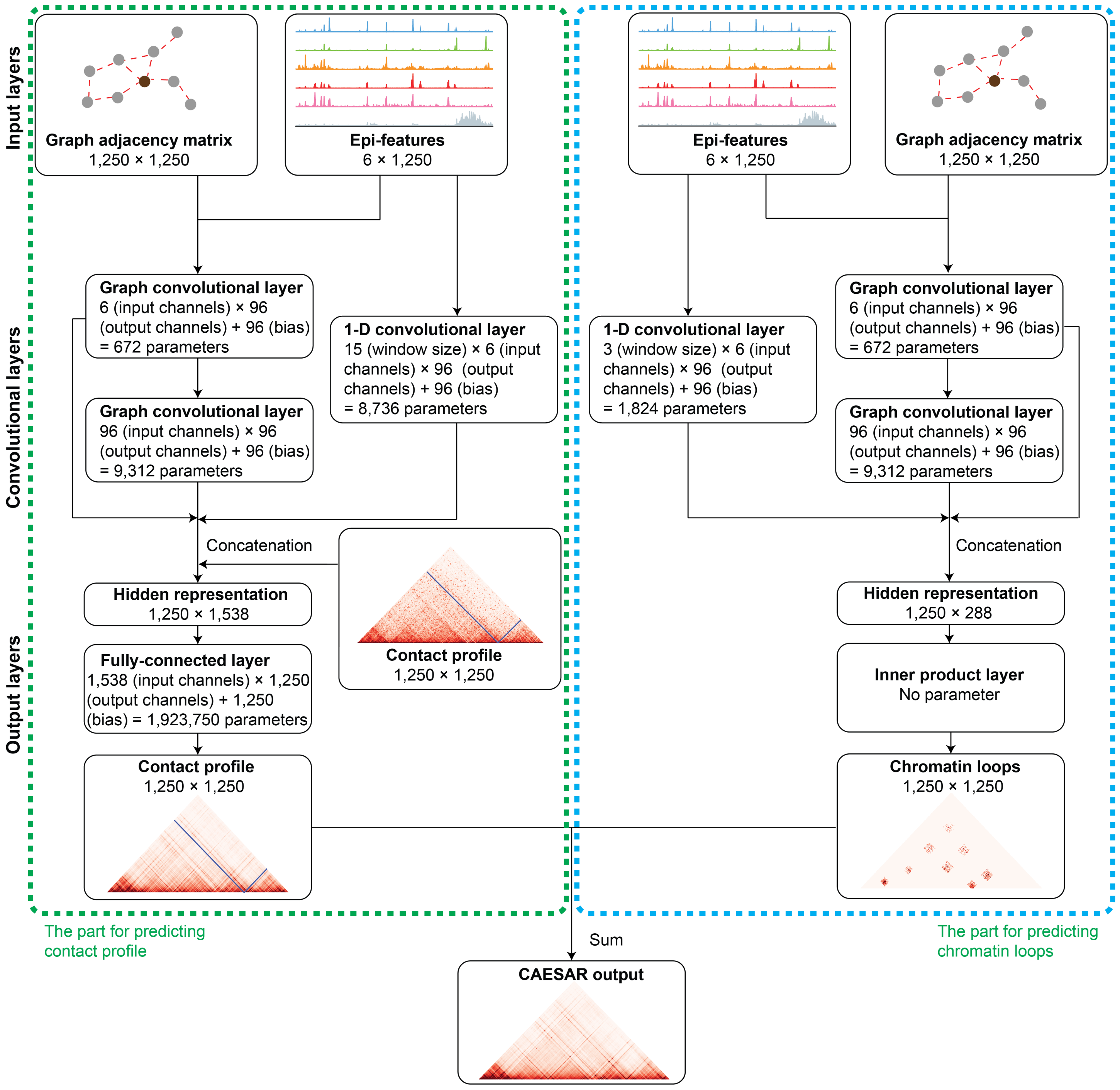


Figure S1: Model structure details.

The model includes two parts — one for predicting chromatin loops and the other for predicting the contact profile, and each part includes input layers, convolutional layers, and output layers. At last, the outputs of the two parts are summed up to generate the final output.

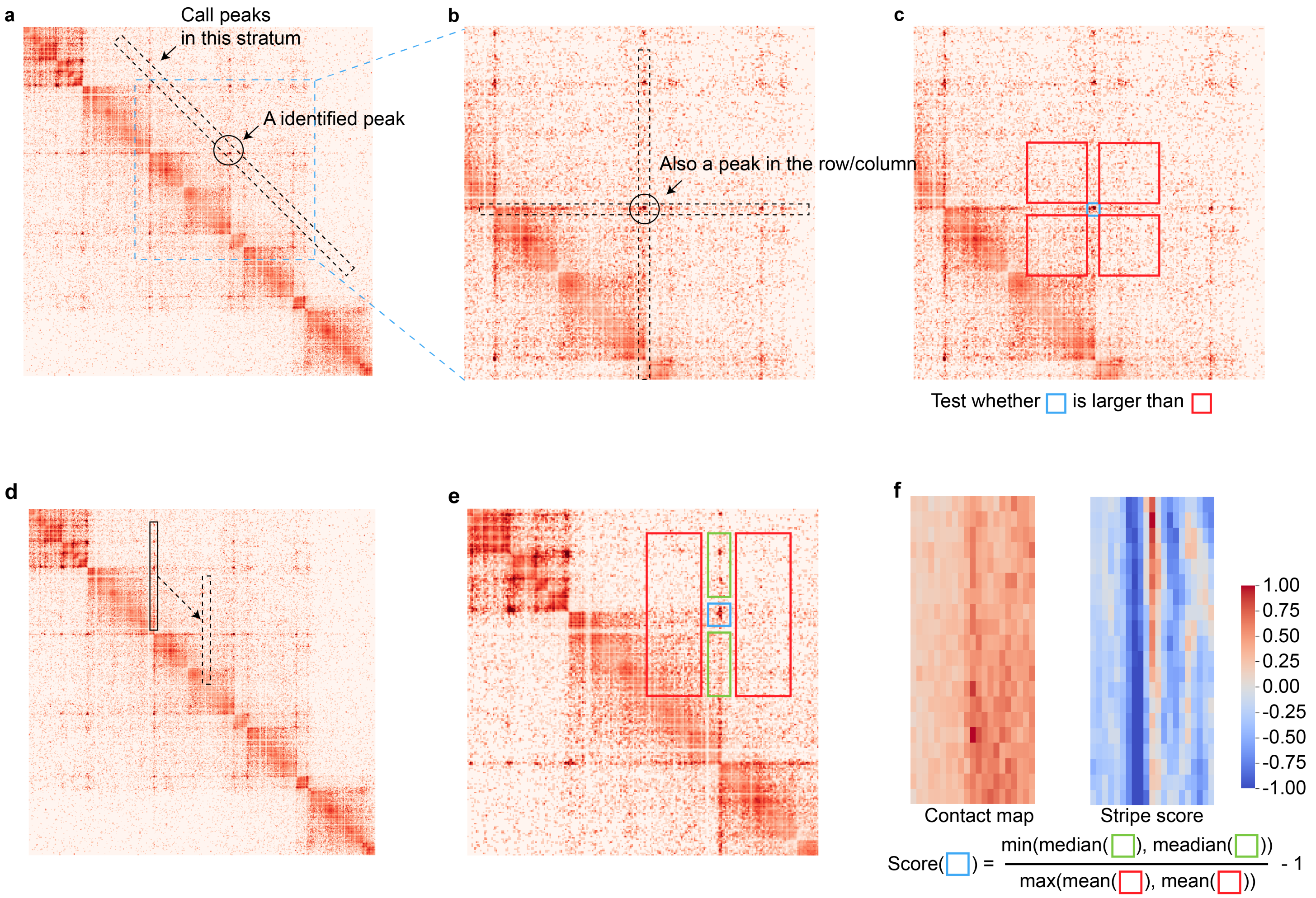


Figure S2: The illustration of the loop caller (a-c) and stripe caller(d-f) in our study.
**a**, Step 1: The peaks on each stratum are called to identify "candidate loops". **b**, Step 2: If a pixel on the contact map is identified as a peak along the stratum, we further evaluate whether it is still a peak on its row and column. **c**, Step 3: Statistical tests (e.g., a *t*-test) are applied to the candidate loops (the blue square), validating whether the value at the pixel is significantly larger than the combination of its immediate neighborhood (red squares). **d**, Step 1: A narrow and long sliding window moves along the diagonal to identify candidate vertical stripes. **e**, Step 2: For each pixel on the candidate stripe, five windows are selected and a "stripe score" is calculated for evaluating whether it is on a stripe. **f**, An example of original contacts versus stripe scores illustrates that positive scores indicate potential stripes.

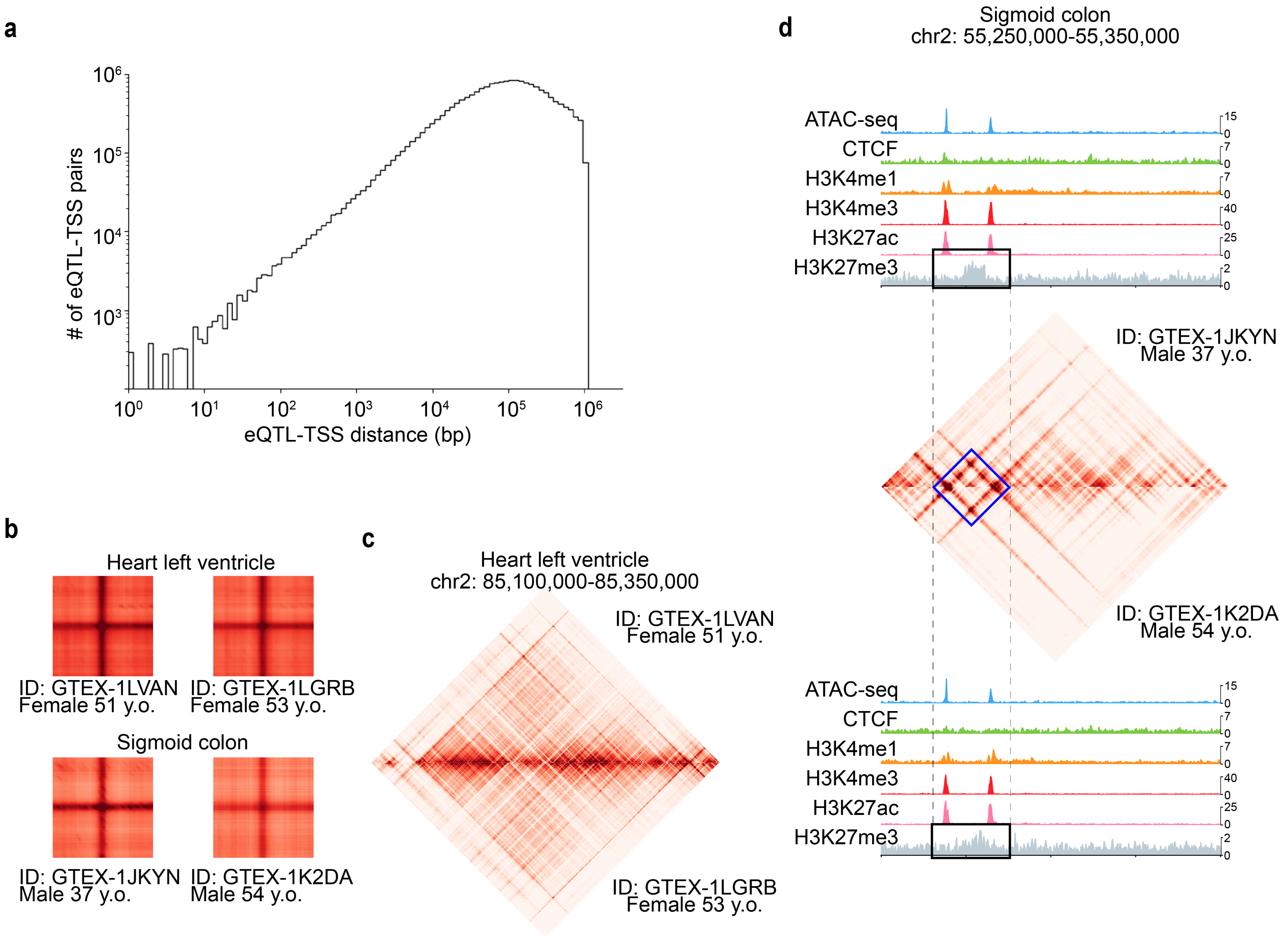


Figure S3: The imputation of high-resolution contact maps and eQTL-TSS enrichment analysis for human tissues. **a**, The distribution of eQTL-TSS distances in the 12 human tissues and cell lines demonstrates that about 50% of eQTL-TSS pairs are less than 100 kb apart, which are hard to identify on low-resolution Hi-C contact maps. **b**, The eQTL-TSS pile-up results for different donors from the heart left ventricle and sigmoid colon are consistent. **c**, The example region from the imputed contact maps of two heart left ventricle donors illustrates that, besides eQTL pile-up results, the imputed tissue contact maps are mostly consistent between individuals. **d**, A counter-example — a loop is observed on the imputed contact map of sigmoid colon from donor GTEX-1JKYN but not donor GTEX-1K2DA, which is related to a more clear H3K27me3 peak in donor GTEX-1JKYN's epigenomic features.

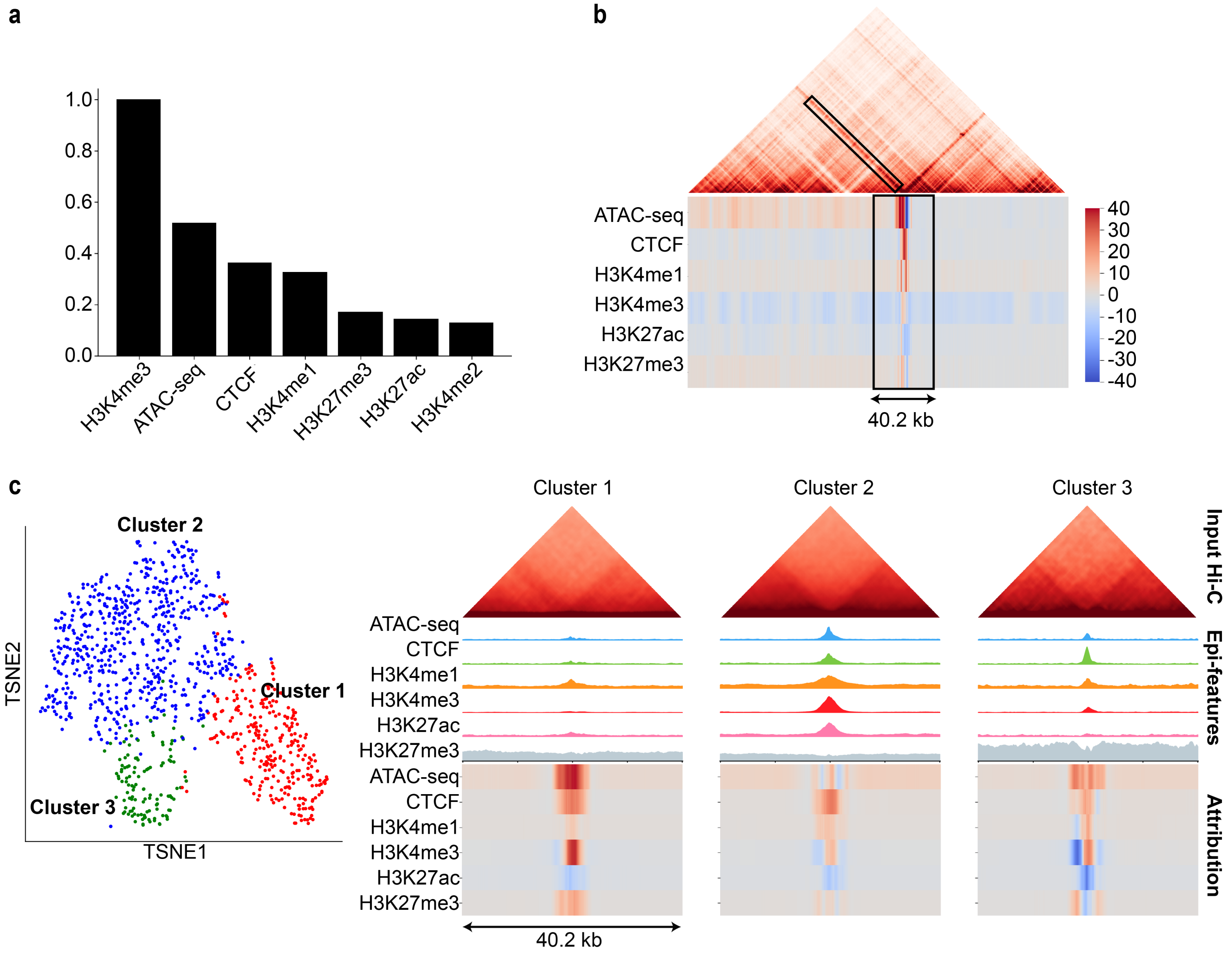


Figure S4: Attributing CAESAR's outputs towards input epigenomic features.
**a**, By attributing the entire contact map to the 7 epigenomic features, we obtained the overall attribution for each epigenomic features. Since H3K4me2 is less commonly profiled and also contributes less, we can leave it out from the 6-epi model. **b**, The attribution is calculated at a stripe region. In the genome-wide attribution analysis of stripes, we collected attribution from the 40.2 kb region centered at the anchor of each stripe. **c**, The clustering and embedding of all stripes' attribution illustrate that there are 3 clusters of stripes, which means the model has learned 3 major patterns indicating stripes. The average input Hi-C, epigenomic features and attribution for each cluster are visualized.

Supplementary Table 1. CAESAR-imputed tissues and cell lines

| Tissue | | |
|---|---|---|
| Adrenal gland | Ascending aorta | Body of pancreas |
| Breast epithelium | Esophagus muscularis mucosa | Esophagus squamous epithelium |
| Gastrocnemius medialis | Gastroesophageal sphincter | Heart left ventricle |
| Lung | Ovary | Pancreas |
| Peyer's patch | Prostate gland | Right atrium auricular region |
| Sigmoid colon | Spleen | Stomach |
| Suprapubic skin | Testis | Thoracic aorta |
| Thyroid gland | Tibial artery | Tibial nerve |
| Transverse colon | Upper lobe of left lung | Uterus |
| Vagina | | |
| **Cell line** | | |
| A549 | A673 | GM12878 |
| GM23338 | HCT116 | HeLa-S3 |
| HepG2 | IMR-90 | K562 |
| Karpas-422 | MCF-7 | MM1S |
| OCI-LY7 | PC-3 | PC-9 |
| SK-N-SH | | |
| **Primary cell** | | |
| B cell | CD14-positive monocyte | Astrocyte |
| Endothelial cell of umbilical vein | Fibroblast of dermis | Fibroblast of lung |
| Foreskin fibroblast | Foreskin keratinocyte | Keratinocyte |
| Mammary epithelial cell | Osteoblast | Skeletal muscle myoblast |
| ***In vitro* differentiated cell** | | |
| Bipolar neuron | Cardiac muscle cell | Hepatocyte |
| Myotube | Neural progenitor cell | Smooth muscle cell |

Supplementary Table 2. Data sources of Hi-C/Micro-C contact maps (with link)

| Contact map | Cell line | 4DN Accession |
|---|---|---|
| Micro-C | H1-hESC | 4DNES21D8SP8 |
| | HFF | 4DNESWST3UBH |
| | mouse ESC | 4DNES14CNC1I |
| Hi-C | H1-hESC | 4DNES2M5JIGV |
| | HFF | 4DNES2R6PUEK |
| | mouse ESC | 4DNESKKSKG7Y |
| | IMR-90 | 4DNES1ZEJNRU |
| | K562 | 4DNESI7DEJTM |
| | GM12878 | 4DNES3JX38V5 |

Supplementary Table 3a. Data sources of epigenomic tracks for human tissues (with link)

| Tissue | Donor* | ATAC-seq** | CTCF | H3K4me1 | H3K4me3 | H3K27ac | H3K27me3 |
|---|---|---|---|---|---|---|---|
| Ascending aorta | F 51 | 422IIZ*** | 846JKO | 202XTW | 645FBM | 982QIF | 103QHX |
| | F 53 | 968TPO | 555DCD | 707AEW | 122LOZ | 069UMW | 589GII |
| Body of pancreas | M 37 | 152PSA | 572DUJ | 827NKO | 876DCP | 520BIM | 977CEC |
| | M 54 | 765MXG | 687APM | 348TQM | 554RQQ | 596PFU | 774CFO |
| Breast epithelium | F 51 | 846ZBX | 661NXJ | 263XKR | 568QQU | 081OTO | 134LLK |
| | F 53 | 654UYP | 304XUZ | 553IAW | 416AUW | 034ZKE | 770WSE |
| Esophagus muscularis mucosa | F 51 | 686ZKE | 443WKD | 701GIE | 403PEI | 894MOX | 200AFX |
| | M 54 | 609GST | 073TPC | 674WSL | 077HGR | 705BTW | 543UBL |
| Esophagus squamous epithelium | F 51 | 096BPX | 266UTR | 658EVN | 773PIU | 204TAU | 049FUB |
| | F 53 | 579BNV | 756URL | 121RSS | 508UPW | 909UAG | 057BFO |
| | M 37 | 944JCE | 559KAB | 525JJM | 621MTP | 522MTS | 188HXK |
| Gastrocnemius medialis | F 51 | 823ZCR | 355ALW | 499VCQ | 098OLN | 601VHO | 453MSI |
| | F 53 | 689SDA | 428BKN | 776EAH | 785DJD | 736ALU | 201OSX |
| | M 37 | 258JCL | 594NSU | 148FWR | 206STN | 801IPH | 519WQH |
| | M 54 | 308HPZ | 998NQG | 161HZJ | 972ETR | 948YYZ | 423LXQ |
| Heart left ventricle | F 51 | 117PYB | 718SDR | 449FRQ | 181ATL | 702OVJ | 613PPL |
| | F 53 | 851EBF | 544APK | 438QZN | 901SIL | 854OXF | 988JLN |
| Peyer's patch | F 51 | 261RWJ | 542SCB | 874HIG | 684EPX | 249IKQ | 491FDG |
| | F 53 | 017RQC | 375VXU | 621BZD | 878KIY | 837SGJ | 982PLJ |
| | M 37 | 954AJK | 419ANE | 416ZMW | 349GPJ | 440PMP | 632SLJ |
| | M 54 | 455GUW | 568IVD | 912XAL | 998QKF | 758KRK | 735VKO |
| Prostate gland | M 37 | 564FZH | 946MNG | 155XVP | 153NDQ | 841AJO | 690CSD |
| Right atrium auricular region | F 51 | 062SVK | 232OFD | 817FGU | 954TSY | 668EVA | 459CKR |
| | F 53 | 984SQJ | 401KRN | 368ORV | 791KFQ | 593KDJ | 793PLF |
| Sigmoid colon | M 37 | 548QCP | 721AHD | 181HTE | 960AAL | 807XUB | 734ZTQ |
| | M 54 | 086OGH | 857RJQ | 775LGE | 172LVU | 937EVN | 860GPM |
| Spleen | F 51 | 078EBD | 595BPR | 831EDZ | 589DBF | 668GBL | 161FEJ |
| | F 53 | 128GBN | 601FEB | 659RJP | 197QDK | 726HTS | 826MTK |
| | M 54 | 850YHJ | 225YGX | 635IRN | 377ILM | 593INW | 080JPX |
| Stomach | F 51 | 641ZPF | 361KVZ | 009RJD | 492BHN | 751BHO | 330MAM |
| | F 53 | 337UIU | 185CCV | 903QBX | 489ZLL | 133NBJ | 357ROS |
| | M 37 | 177NIJ | 618QYE | 493MQY | 843UEZ | 944KAZ | 227DGG |
| Testis | M 37 | 210NKB | 753RME | 956VQB | 611DJQ | 136ZQZ | 503QSX |
| Thyroid gland | F 51 | 201FIW | 955BIB | 497OVD | 309UVT | 500YBS | 586DVD |
| | M 37 | 749MUH | 505ZGX | 906YES | 901BRV | 597BWL | 748LUA |
| | M 54 | 474XFV | 033KMZ | 639NMN | 975NOU | 203KCB | 582PKH |
| Tibial artery | M 37 | 102RSU | 079YAP | 960VRR | 780CNW | 891BTJ | 764OHK |
| Tibial nerve | F 51 | 831KAH | 793YAD | 338PGG | 677MOE | 778QHG | 992XOO |
| | F 53 | 100TUY | 875NEW | 850RVA | 314SPW | 771YJT | 611YUJ |
| | M 37 | 484UAU | 434XLP | 981CTV | 384MUF | 516LQO | 860ZCZ |
| | M 54 | 508FVM | 689VEF | 590NNJ | 464TRM | 091KXI | 662ASZ |
| Transverse colon | F 51 | 386HAZ | 449SEF | 500QVK | 315EZG | 792VLP | 604QMH |
| | F 53 | 404LLJ | 236YGF | 791LZY | 933BVL | 208QRN | 840VWD |
| | M 37 | 668VCT | 608WPS | 516QFO | 813ZEY | 640XRV | 643KID |

| | | | | | | | |
|---|---|---|---|---|---|---|---|
| Upper lobe of left lung | F 51 | 323UTX | 799TJD | 238WIK | 429VWL | 453MUW | 706OFD |
| | F 53 | 702DPD | 224WWI | 263OXW | 208WDY | 738SXD | 859MXQ |
| | M 37 | 164WOF | 027FSZ | 595MTV | 074WIB | 505YFA | 469YCE |
| | M 54 | 650FLQ | 463XCZ | 348FGT | 701FGA | 948TOS | 050LBS |
| Uterus | F 53 | 129BZE | 684PGO | 035ONO | 354ZUG | 249INE | 111DTF |
| Vagina | F 51 | 733YNW | 655ECZ | 495RJG | 647HAQ | 346FVK | 278TQE |
| Adrenal gland | mixed | 277KRY | 899JSO | 455JUO | 620TXL | 094VJC | 181JFC |
| Gastroesophageal sphincter | mixed | 260ZIV | 298ZPF | 134KZX | 037GFN | 600TOW | 965BLU |
| Lung | mixed | 647AOY | 000DMH | 356ANC | 466DZW | 540ADS | 204NFO |
| Ovary | mixed | 712PYJ | 548DDS | 113AFY | 139TLA | 268JQE | 037SNV |
| Pancreas | mixed | 595HZQ | 000DND | 984UHU | 315LPR | 402HFW | 486NDF |
| Suprapubic skin | mixed | 709IYR | 485VQV | 374XIN | 362QYU | 413QLR | 410BWN |
| Thoracic aorta | mixed | 344ZTM | 549TXG | 803IBD | 930HLX | 318HUC | 939RLS |

* In this column, “F” and ”M” indicate female and male, and numbers indicate the donors’ age. “Mixed” indicates the datasets are from multiple donors.

** For tissues or cell lines without available ATAC-seq data, we collected DNase-seq instead.

* * * “422IIZ” is short for “ENCSR422IIZ”. In this and the following tables, “ENCSR” is omitted for all accessions.

Supplementary Table 3b. Data sources of epigenomic tracks for human cell lines (with link)

| Cell line | DNase-seq | CTCF | H3K4me1 | H3K4me3 | H3K27ac | H3K27me3 |
|---|---|---|---|---|---|---|
| A549 | 000ELW | 000DNA | 636PIN | 000DPD | 778NQS | 000AUJ |
| A673 | 346JWH | 611JJS | 521IZK | 435FGK | 714TJD | 747BYL |
| GM12878 | 000EJD | 000DZN | 000AKF | 057BWO | 000AKC | 000DRX |
| GM23338 | 004SUL | 987GXT | 249YGG | 657DYL | 729ENO | 386RIJ |
| HCT116 | 000ENM | 240PRQ | 161MXP | 333OPW | 661KMA | 810BDB |
| HeLa-S3 | 959ZXU | 000DLO | 000APW | 340WQU | 000AOC | 000APB |
| HepG2 | 149XIL | 000DUG | 000APV | 575RRX | 000AMO | 000AOL |
| IMR-90* | 477RTP | 000EFI | 831JSP | 087PFU | 002YRE | 431UUY |
| K562 | 000EKS | 000DWE | 000EWC | 668LDD | 000AKP | 000EWB |
| Karpas-422 | 019JDO | 113REG | 306VSH | 910XKX | 660IQS | 963HAR |
| MCF-7 | 000EPJ | 560BUE | 493NBY | 985MIB | 752UOD | 761DLU |
| MM1S | 458LIB | 402IDP | 094VCE | 361FWQ | 758OEC | 404LJZ |
| OCI-LY7 | 489NAM | 027HML | 060WGK | 005SXO | 447ZGY | 752KQT |
| PC-3 | 052AWE | 359LOD | 566UMF | 275NCH | 826UTD | 881TWJ |
| PC-9 | 940NLN | 243INX | 913MGR | 441JWF | 769FOC | 726LZG |
| SK-N-SH | 000EPZ | 541AMF | 661BMA | 975GZA | 564IGJ | 914QOK |

* IMR-90 is not a cancer cell line.

Supplementary Table 3c. Data sources of epigenomic tracks for primary cells (with link)

| Primary cell | DNase-seq | CTCF | H3K4me1 | H3K4me3 | H3K27ac | H3K27me3 |
|---|---|---|---|---|---|---|
| B cell | 381PXW | 000AUV | 290YLQ | 000DQR | 000AUP | 162DGX |
| CD14-positive monocyte | 000EPK | 000ATN | 000ASM | 000DWL | 000ASJ | 000DWM |
| Astrocyte | 000EPM | 000AOO | 000AOT | 000AOU | 000AOQ | 000AOR |
| Endothelial cell of umbilical vein | 000EOQ | 000ALA | 000AKL | 578QSO | 000ALB | 000AKK |
| Fibroblast of dermis | 000EPO | 000APM | 000ARV | 000APR | 000APN | 000APO |
| Fibroblast of lung | 000EPR | 000DWY | 000AMU | 915QOL | 000AMR | 000AMS |
| Foreskin fibroblast | 153LHP | 000DUH | 367HVD | 813CFB | 917QEH | 417IEJ |

| | | | | | | |
|---|---|---|---|---|---|---|
| Foreskin keratinocyte | 035RVH | 817HTJ | 027BAJ | 075OQB | 666TFS | 377MRR |
| Keratinocyte | 000ELH | 000DNC | 000ALI | 970FPM | 000ALK | 000DWU |
| Mammary epithelial cell | 000ENV | 000DUS | 521FND | 000DUQ | 000ALW | 000ALX |
| Osteoblast | 000ELJ | 000APF | 000APJ | 000ATH | 000APH | 000AQS |
| Skeletal muscle myoblast | 000EOO | 000ANE | 000ANI | 596NOF | 000ANF | 000ANG |

Supplementary Table 3d. Data sources of epigenomic tracks for *in vitro* differentiated cell (with link)

| *In vitro* differentiated cell | DNase-seq | CTCF | H3K4me1 | H3K4me3 | H3K27ac | H3K27me3 |
|---|---|---|---|---|---|---|
| Bipolar neuron | 626RVD | 619IUE | 301AEA | 849YFO | 905TYC | 472SEY |
| Cardiac muscle cell | 842KCP | 713SXF | 276OLB | 652QNW | 000NPF | 864LRY |
| Hepatocyte | 364MFN | 252QYR | 689QUB | 442ZOI | 507UDH | 637RLN |
| Myotube | 000EOP | 000ANS | 000ANX | 000ANZ | 000ANV | 000ATI |
| Neural progenitor cell | 963ALV | 125NBL | 274OIJ | 661MUS | 449AXO | 139PIA |
| Smooth muscle cell | 248CME | 261VAS | 130IMV | 515PKY | 210ZPC | 143RMH |

### 1 Data collection and processing

The datasets used in our cross-validation experiments include Hi-C contact maps, epigenomic features, and Micro-C contact maps for three cell lines — hESC, mESC and HFF. Micro-C and Hi-C contact maps of HFF, hESC, and mESC were downloaded from the 4DN data portal [1]. Chromatin accessibility (ATAC-seq and DNase-seq) data of HFF, hESC, and mESC were downloaded from ENCODE database [2]. Twelve ChIP-seq signals of mESC and hESC (CTCF, Rad21, Nanog, H3K4me1, H3K4me2, H3K4me3, H3K9ac, H3K9me3, H3K27ac, H3K27me3, H3K36me3, and H3K79me2) were downloaded from ENCODE database and CistromeDB [3, 4]. Due to the unavailability of HFF ChIP-seq data, we used CUT&RUN as an alternative, and six HFF CUT&RUN signals (CTCF, H3K4me1, H3K4me2, H3K4me3, H3K27ac, and H3K27me3) were downloaded from the 4DN data portal[1].

For imputing high-resolution contact maps of additional human tissue types and cell lines, we collected Hi-C contact maps and epigenomic signals of human tissues and cell lines. Hi-C contact maps with more than 1 billion contacts (K562, IMR-90, and GM12878) were downloaded from the 4DN data portal. The epigenomic signals from 91 samples were downloaded from ENCODE (Supplementary Note 2).

To validate CAESAR's performance in predicting the interactions between regulatory elements, we collected eQTLs and CRISPRi data. The eQTL data of 10 human tissues (adrenal gland, sigmoid colon, transverse colon, heart left ventricle, lung, tibial nerve, ovary, pancreas, spleen, and stomach) and 2 human cell lines (GM12878 and IMR-90) were downloaded from GTEx Analysis Release V8 [5]. The K562 CRISPRi data were downloaded from the original study of Fulco *et. al.* [6].

To evaluate CAESAR's performance in different genomic regions, we collected phastCons scores and repli-seq data to separate all regions into different groups. The 100-way phastCons scores of hg38 were downloaded from UCSC genome browser [7], and the repli-seq data were downloaded from the 4DN data portal.

The detailed metadata is summarized in Supplementary Tables 2 and 3. All Hi-C contact maps were processed into 1 kb resolution and then linearly interpolated to 200 bp resolution; all Micro-C contact maps and epigenomic signals were processed into 200 bp resolution. All Micro-C contact maps were OE-normalized (i.e., observed/expected normalized for each stratum). All mouse data in our analysis used mm10 reference genome, and all human data in our analysis used hg38 reference genome.

### 2 Collecting epigenomic signals from ENCODE database

We searched the ENCODE Data Matrix (https://www.encodeproject.org/matrix/?type=Experiment) to collect epigenomic signals for imputing high-resolution contact maps. We limited the organism to *homo sapiens*, and identified all biosamples (tissues, cell lines, primary cells, and *in vitro* differentiated cells) with all of the following signals — ATAC-seq/DNase-seq, CTCF, H3K4me1, H3K4me3, H3K27ac, and H3K27me3. For a specific human tissue, there are two outcomes. If all six epigenomic tracks are available for individual donors, then we will impute the contact map for these individual donors separately. If we do not have sufficient epigenomic tracks for imputing for the individual donors, then we will only impute one contact map for the specific human tissue (referred to as "mixed-donor tissue"). In the end, we identified 91 sets of epigenomic signals from 50 individual donors for 21 tissue types, 7 mixed-donor tissues, 16 cell lines, 12 primary cells, and 6 *in vitro* differentiated cells.

### 3 Detailed CAESAR model structures

The model includes two major parts — one for predicting chromatin loops, and the other for predicting contact profile. Each part includes consecutive input layers, convolutional layers, and output layers (Figure S1). CAESAR captures the interpolated Hi-C contact map as a graph $\mathcal{G}$ with nodes representing genomic regions of 200 bp long, and weighted edges representing chromatin contacts. $A$ is the adjacency matrix of $\mathcal{G}$. For both parts, the inputs include the graph adjacency matrix $A$ and the epigenomic features $X$. As one 250 kb region is fed into the model each time, the dimension of the input adjacency matrix is 1250×1250. In a 6-epigenomic model, the size of the feature matrix is 6×1250.

In deep learning models, convolutional kernels are small filters sliding through the input to extract certain patterns. When the filter is applied to an input element, it calculates the weighted sum of the element with its local neighbors. In a convolutional layer, multiple kernels work in parallel to learn different sets of weights and extract different patterns. There are two types of convolutional layers, 1-D convolutional (Conv1D) and graph convolutional (GC) layers in CAESAR. Conv1D layers operate along the genome fiber, aggregating the epigenomic features from nearby bins. GC layers extract spatial epigenomic patterns over the spatial neighborhood specified by $\mathcal{G}$. Here, we use the GC layer

$$Y = \sigma(\tilde{A}XW)$$

in which $X$ and $Y$ are the input and output, $\tilde{A}$ is the normalized graph adjacency matrix, $W$ is the trainable parameters, and $\sigma$ is the $relu$ activation function [8]. GC layers provide additional structural patterns for imputing high-resolution chromatin architecture. For example, if two distant loci $i$ and $j$ are in the same TAD, then nodes $i$ and $j$ are neighbors on the graph. Therefore, when we predict the contact profile of $i$, the information flows from $j$ to $i$ in the GC layers, so that the features at $j$ contribute to the prediction of $i$, and *vice versa*. The window size for each 1-D convolution kernel is 15 in the contact profile predicting part and 3 in the loop predicting part, which captures relevant features from a 3kb and 600 bp neighborhood, respectively.

For the contact profile predicting part, the output layer is a fully-connected layer. The input of this layer is the concatenation of convolutional layers' outputs and the Hi-C contact profile, and the output is the imputed contact profile of each 200 bp bin. For the loop predicting part, the output layer is an inner product layer. This layer also takes the concatenation of convolutional layers' outputs as input, and calculates the inner product between each bin pairs' representation to predict the chromatin loops. The outputs of the two output layers are summed up to generate the final imputation result. The model includes 2 million parameters, which is much fewer than the number of elements (∼15 billion) in the contact matrix.

### 4 Train, test and tune sets of chromosomes

We split the chromosomes into three sets of comparable sizes to train, tune, and test our CAESAR model. For hg38, the train set include chr1, 4, 7, 10, 13, 17, and 18 (total length 1,010,309,426 bp), the test set include chr2, 5, 8, 11, 14, 15, 21, and 22 (total length 1,010,520,404 bp), and the tune set include chr3, 6, 9, 12, 16, 19, 20, and X (total length 1,010,212,587 bp). For mm10, the train set include chr1, 4, 7, 8, 10, and 11 (total length 879,600,295 bp), the test set include chr2, 5, 9, 12, 14, and X (total length 874,605,583 bp), and the tune set include chr3, 6, 9, 13, 15, 16, 17, 18, and 19 (total length 879,570,794 bp).

### 5 Hyperparameter Tuning

CAESAR includes two hyperparameters: 1) the number of convolutional layers and 2) the number of convolutional kernels in each convolutional layer. We examine 4 different convolutional layer configurations: i) 3 GC layers, ii) 2 GC layers and 1 Conv1D layer, iii) 1 GC layer and 2 Conv1D layers, and iv) 3 Conv1D layers. In each layer, we tested 3 different numbers of convolutional kernels - 64, 96, and 128.

For each of the 12 combinations, we trained a CAESAR model with the train set and evaluated with the mean squared error (MSE) on the tune set, and the model with 2 GC layers, 1 Conv-1D layer, and 96 kernels at each layer, achieved the best performance.

### 6 Baselines methods and parameters

In existing literature, there are two major categories of machine learning approaches for imputing Hi-C contact maps. The first category takes low-resolution contact maps as input and treats Hi-C contact maps as 2-D images, exemplified by HiCPlus [9]. The second category predicts the contacts between every two bins with genomic or epigenomic features from the two bins, exemplified by HiC-Reg [10]. Therefore, we use HiCPlus and HiC-Reg as two baselines in our experiments.

HiCPlus is a deep-learning model with three sequential layers, in which the first and third layers are Conv2D layers, and the second layer is a fully-connected layer. Since the matrices in our study are much bigger, we accordingly increased both the number and the size of Conv2D kernels. We set the number of Conv2D kernels to be 96, and the size of Conv2D kernels to be 15×15. The model was re-trained with hESC train set, in which the inputs were the Hi-C contact maps and the targets were the Micro-C contact maps.

HiC-Reg uses random forests (RF) to predict the contacts between locus $i$ and $j$ with the epigenomic features near $i$ and $j$ as well as the distance between $i$ and $j$. We used a 240-tree RF to re-train the model with hESC train set, in which the combination of 6 epigenomic features were the input and the Micro-C contact maps were the targets.

### 7 Fast loop calling

HICCUPS [11] was designed to call loops from Hi-C contact maps at lower resolutions, and we observed that it failed to call loops on our imputed contact maps even if the loop can be easily observed. The reason may include 1) our imputed contact maps are OE-normalized rather than KR-normalized, and 2) only short-range interactions are imputed by CAESAR — both of which are unfavorable for HICCUPS to calculate the statistical significance of chromatin loops. Therefore, we implemented a fast loop calling approach. This approach has two steps: peak calling and enrichment score calculation. First, we take the strata as the inputs and call peaks on each stratum with Python "scipy.find_peaks" function (Figure S2a).

If a pixel on the contact map is identified as a peak on the stratum, we further check whether it is still a peak on its row and column (Figure S2b). Only elements that peak on all 3 directions are labeled as candidate loops. The peak calling accelerates loop calling by quickly narrowing down the searching range. Afterward, *t*-test is applied to the candidate loops, validating whether the value of the pixel is significantly larger than the combination of its surrounding regions (Figure S2c). The threshold we used in this step was 0.005. For the differential analysis between two contact maps, we say two loops "match" if they are less than 2 kb apart.

### 8 Stripe calling with Quagga

We developed an original method, Quagga, to call stripes on the CAESAR-imputed, Micro-C, and Hi-C contact maps. Stripes are labeled as "vertical" or "horizontal" according to their directions on the top right half of the contact map. Quagga calls vertical and horizontal stripes separately. Quagga identifies vertical stripes as follows. First, a narrow, long sliding window anchored at the diagonal moves along the diagonal of the OE-normalized contact map. The contacts are summed up in the window at each step to obtain a 1D vector, and the peaks of the vector are identified as the candidate vertical stripes (Figure S2d). In our work, we used a 100×1 sliding window at 1 kb resolution. However, loops, TAD boundaries, or random noise can be false positives, and therefore we further calculate a "stripe score" (Figures S2e and S2f). For each pixel on a candidate stripe, five regions are chosen: a $i \times i$ square $X$ centered at it, two neighboring $j \times i$ regions along the candidate stripe (upper: $X_u$; lower: $X_w$), two $(2j+i) \times j$ regions on the left ($X_l$) and right ($X_r$) and the "stripe score" is calculated as

$$Score = min(median(X_u), median(X_w))/max(mean(X_l), mean(X_r)) - 1.$$

In our work, we set $i$=1 and $j$=10. The left and right regions work as the background, and taking the maximum of the two avoids TAD boundaries to be falsely called as stripes. Calculating the median of the upper/lower regions ensures a single large value (e.g., a loop) does not increase the score. If a pixel is on a vertical stripe, then the enrichment score should be greater than 0. At last, Quagga calculates the summation of the stripe scores for each candidate stripe and output the anchor position if the summation is above a threshold. For the differential analysis between two contact maps, we say two stripes "match" if their anchor positions are less than 2 kb apart.

When we applied Quagga to the contact map imputed with 3 epigenomic features (ATAC-seq, CTCF, and H3K27ac), more than 20,000 stripes were called. Since CAESAR outputs each 200 bp bin's contact profile separately, when the input does not provide sufficient information about chromatin structures, the model outputs from neighboring bins are more random and inconsistent. The inconsistency of rows/columns may result in the calling of false-positive stripes, which explains the over-prediction of stripes by the 3-epi model.

### 9 The genome-wide attribution analysis of stripes

The integrated gradient can be applied to arbitrary regions of the imputed contact map. Here we show, by calculating the attribution of all stripe regions, we can identify sub-types of stripes.

We selected the stripes which were called on both Micro-C and CAESAR-imputed contact maps. Since the stripes were called at 1 kb resolution, we identified the accurate stripe anchors at 200 bp resolution by selecting the row/column with the largest summation on the Micro-C contact map. The stripe regions were defined as 11×500 long rectangles starting from the stripe anchor on the diagonal and stretching in the same direction as the stripes. We calculated the attribution of all stripe regions with integrated gradient. Only the attribution near the anchors (i.e., 100 bins both upstream and downstream) were preserved (Figure S5b), resulting in 6×201 attribution matrices, which were flattened to 1,206-dim vectors. Afterwards, we used PCA to reduce the dimension from 1,206 to 50, and then *k*-means to cluster the 50-dim vectors. The 50-dim vectors are further transformed into 2-dim with *t*SNE for visualization.

The attribution near all stripe anchors can be clustered into 3 groups, in which each group has its characteristic patterns in average attribution, input Hi-C contact maps, and epigenomic features (Figure S5c). Cluster 1 has fewer Hi-C contacts, lower epigenomic peaks, and positive attribution of ATAC-seq, CTCF, and H3K4me3/H3K27me3, which demonstrates that in regions with less enriched contacts and epigenomic features, the peaks of ATAC-seq, CTCF, and H3K4me3/K27me3 tend to be connected to stripes by CAESAR. Cluster 2 demonstrates that, at TAD boundaries, the peaks of CTCF and H3K4me1/H3K4me3 closely relates to stripes. The characteristic pattern of cluster 3 is the negative H3K27me3 attribution at the stripe anchor and positive attribution surrounding it, which means CAESAR tends to identify the loci next to H3K27me3-enriched regions as stripe anchors. Although the sub-types still need to be further explored and experimentally validated, this approach provides interpretable insights into our "black box".

### 10 Web server implementation

The imputed high-resolution contact maps are shared on a web server (https://nucleome.dcmb.med.umich.edu/), which allows users to easily navigate these fine-scale chromatin structures, and the corresponding explanatory epigenomic features. The back-end of the server uses python *Flask* with *sqlite*. The front-end of the server uses *bootstrap* framework. The web server utilizes multi-threading to allow multiple users to access it at the same time. Our web server processes host data at multiple ports at localhost. We use *Nginx* to perform the reverse proxy that passes internet requests to them. After contact maps are generated, we run *Nucleome Browser* on our web server. Nucleome Browser is an open platform to integratively and interactively browse coordinate-based genome data. Nucleome Browser extends conventional track-based genome browsing to panel-based genome browsing, thus breaks the linear limitation of stacked tracks view mode. Different panel modules host and render different modality data including visualized tracks and reconstructed 3D chromatin structures.
